# Spatial Modeling of Tissues for Morphogenic Field Analysis

**DOI:** 10.64898/2025.12.06.692715

**Authors:** Aaron Osgood-Zimmerman, Micha Sam Brickman Raredon

## Abstract

Tissues are shaped by extracellular signaling fields which convey information between cells. The cellular composition of tissues, and the extracellular signaling within the tissue, are innately spatially structured. Modern spatialomics data provide unprecedented measurement of ligand and receptor expressivity *in situ* from tissue sections. Here, we show that by adapting generalizable geospatial statistical models to spatialomics data, we are able to reveal statistically-detailed portraits of morphogenic field interactions within tissues and thereby approach a richer set of biologic questions than is typically pursued. The general methods piloted here can readily be applied to spatialomics data from diverse platforms with no need to alter data collection techniques. Our results demonstrate that the application of spatial statistical modeling to spatialomics data opens many avenues for future experimentation that will be valuable to fundamental biology and to regenerative medicine.

**Graphical Abstract:** 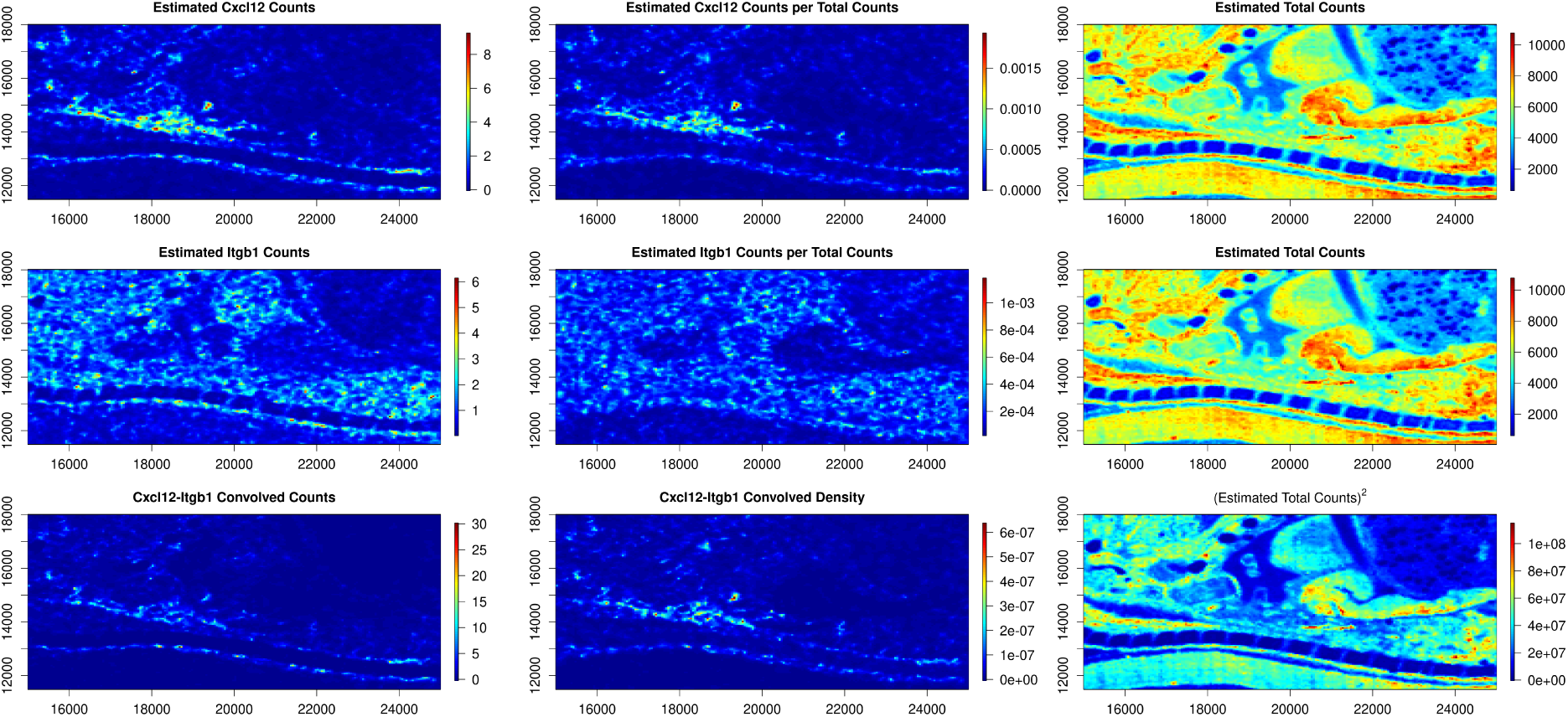

**Highlights:**

- Tissue biology & regenerative medicine requires analysis of tissue morphogen fields and morphogenic interactions
- Spatial statistics can be used to model continuous morphogenic interaction fields in tissues from discrete spatialomics data

## 1. Introduction

### 1.1. Morphogenic Field Interactions are Fundamental to Tissue Biology and Regenerative Medicine

Cells within tissues are continuously processing information and updating their states, receiving cues from their local microenvironment and contributing cues to their local milieu. The study of cellular connectivity (the potential for information flow between cells) and cell-to-cell signaling (information flow between cells) is fundamental to the study of tissue biology. Accurate modeling of cellular connectivity and cell-to-cell signaling, particularly through ligand-receptor syntax, provides a quantitative foundation for understanding tissue-scale changes and dynamics.

It has long been understood that tissues are shaped by morphogenic signaling between cells (*1–4*). During development, homeostasis, disease, and regenerative remodeling, cells within tissues release signaling molecules which diffuse through the extracellular matrix and can be locally received and processed by resident cells. Until recently, it has only been possible to capture data on a small number of morphogenic mechanisms at once from a single tissue. However, it is the *holistic set* of soluble and insoluble cue intensities – taken together as a single combinatoric ‘input’ (*5–7*)– that encodes the full spectrum of locally-transducable signaling information, often irreducible to the individual cues themselves (*8–10*). In other words, extracellular signaling is intrinsically high-dimensional — information is encoded via specific combinations of cues at particular levels which are processed in parallel by resident cells — and is intrinsically spatially patterned: as an organism develops, intersecting 3-dimensional signaling gradients combine to drive tightly localized cellular differentiation, proliferation, and migration events. These morphogenic field interactions are major governors of tissue-scale emergent behavior including developmental morphogenesis and adult tissue inflammation & homeostasis (*11–14*). To understand tissue-scale change, therefore, we need robust tools to study high-dimensional signaling and connectivity dynamics in both time and space.

Spatialomics is newly presenting scientists with an opportunity to estimate thousands of morphogenic fields at once, at both the raw and relative information level, thereby allowing us to model and visualize the specific complex patterns of extracellular signaling that controls cellular differentiation and transition in living tissues. However, to properly handle and analyze these data, we need appropriately designed statistical approaches. This paper presents one such approach, and shows that by adapting techniques common in geospatial data analysis, we are able to newly model and visualize the high-dimensional patterns of extracellular signaling that have long been theorized to be fundamental to tissue growth and change.

### 1.2. The Critical Need for Spatial Statistics in Tissue Biology

Spatialomics data differ from the true state of the tissue in two important ways. First, regardless of platform resolution, all spatialomics data collection methods collect data at discrete locations, necessarily creating error in reported spatial sampling coordinates. Second, no bulk RNA, single cell, or spatialomics methods perform true censuses – that is, they do not measure every single molecule within every sample, and in many cases measure only a small fraction of the total present molecules (*15, 16*). As a result, observed data from all spatial transcriptomic methods is subject to sampling variability. Ignoring sampling variability while performing data-analysis can lead to bias, over- or under-confidence in results, and false findings (*17*). Properly accounting for the sampling variability in any dataset requires a statistical model aligned with the data-generating mechanism.

Our statistical goal in this work was to take spatialomics data observations as input and to produce output which estimates the true state of the tissue, acknowledging and accounting for the important ways in which the data differs from the tissue itself. That is, we need statistical models to take transcriptomic data measured at **discrete** spatial locations as input and to produce estimates of **continuous** spatial fields as outputs, along with estimates of uncertainty, which act as a statistical prediction of transcriptomic counts across the spatial observation domain. Spatial statistics has a rich literature of tools to leverage which are designed exactly for this purpose. Furthermore, they account for the spatial autocorrelation present within these datasets, which, if ignored, can again lead to incorrect uncertainty estimates and false findings (*18*).

### 1.3. Biological Tissues are Spatially Structured

Advances in genetics data collection methodologies have led to rich, beautiful datasets which can easily be mistaken to be “the truth.” Spatialomics, loosely defined as the group of technologies capable of measuring molecular characteristics of tissues within their natural spatial alignment, has justifiably been heralded as a transformative frontier in life sciences. However, the quickly developing methods, and the immense amounts of data produced, have outpaced *analysis techniques*, which need to mature (*19*).

Spatialomics collection methods have been developed precisely because there is observable and necessary spatial structure apparent in tissues. Yet, this fact isn’t often explicitly incorporated in analyses. The spatial structure of healthy vs. diseased tissues is often radically dissimilar, but many currently-practiced computational methods are descendants of single cell techniques which were necessarily non-spatial. Ignoring the spatial organization, at best, may be omitting useful information and, at worst, may be lead to an incorrect interpretation of the data. When approaching morphogenic field analysis, i.e., the quantitative study of high-dimensional extracellular signaling mediated by ligand-receptor binding interactions via the extracellular milieu, it is essential to take these principles into account, or we risk misinterpreting statistically spurious findings.

In this manuscript we introduce a flexible continuous spatial modeling framework that can be used to estimate one or more features measured via spatialomics. The proposed approach also provides uncertainty estimates of each quantity over space, which allows propagation of uncertainty to downstream outputs (such as ligand-receptor binding density, the main biologic focus of this paper.) The modeling in this manuscript is meant to be a launchpad for the use of powerful, existing frameworks to learn more readily and deeply from spatialomics data, thereby decreasing the chance of generating unreproducible results. We use our bespoke pipeline to quantitatively assess a portion of a public dataset capturing mammalian embryonic organogenesis. We demonstrate via our results that complex domain architecture in tissues can be derived directly from spatial ligand-receptor connectivity patterns, even without study of transcripts coding for intracellular proteins.

## 2. Methods

### 2.1. Overview

#### 2.1.1. Principles of Ligand-Receptor Analysis

Our focus in this paper was to model a useful proxy for local ligand-receptor binding *density*. This is different from local ligand-receptor connectivity *character*, which is the primary focus of many cell-to-cell communication inference tools including our prior work (*20*). This distinction is non-trivial. When ligand-receptor connectivity *character* is computed, individual feature values represent a proxy for *relative* signal strength; when ligand-receptor binding *density* is computed, individual feature values represent a proxy for *absolute* signal strength. Our focus on binding density in this paper, rather than character, allows us to propagate our outputs to downstream models more easily and with fewer artifacts, as is shown in the Results section. By applying spatial statistics to the raw counts in spatial transcriptomics, rather than the normalized counts, our model outputs are easily treated as a new fundamental layer of information for multi-modal data analysis, one that directly estimates local ligand-receptor binding density within the tissue. This output can then be normalized, similarly to how we usually normalize local gene counts to relative values, to yield proxies for local ligand-receptor connectivity *character* more familiar to most in the field.

All connectivity tools require us to convolve information from the ligand field with information from the receptor field. Historically, this has been done by computing either the product (*20*) or the geometric mean (*21*) of all subunits involved. Here, we have experimentally pioneered the use of alternative approaches to convolving the ligand and the receptor values. Assuming that the counts of individual transcripts are proportional to the counts of individual proteins (an almost certainly *incorrect* assumption, but one that we are forced to accept at this time) we may treat ligand-receptor binding as a stoichiometric process, i.e., a function of the concentration of the ligand and the concentration of the receptor.

Ligands diffuse locally, with a characteristic diffusion distance roughly proportional to the molecular weight and chemical character of the molecular species and how it interacts with other matrix elements. Extracellular ligands can be sensed and processed by tissue-resident cells expressing specific receptor profiles. All previous computational approaches address this fundamental property of tissue biology by first creating local neighborhoods in histologic space and then computing cell-to-cell ligand-receptor connectivity within each neighborhood. When working with continuous field models, as is demonstrated here, local neighborhood construction is no longer necessary, because the spatial field, intrinsically, can be considered as a filter which takes in noisy spatial data and returns a smoothed version. Although the smoothed output is technically an estimate of the underlying state of the tissue, in practice the smoothing process can be viewed as sharing locally-measured information between nearby areas. This modeling approach thereby builds a robust framework for convolution of ligand and receptor fields. The fields themselves can be independently tuned to reflect specific molecular properties, and the convolution approach customized to reflect binding kinetics.

#### 2.1.2. Principles of Spatial Modeling

Spatial statistics is a field of statistics used to study datasets in which the location of the data observations is recorded. Typically, a spatial statistical model will explicitly account for spatial dependence among observations to acknowledge that spatially nearby observations are often more strongly correlated than observations that are located further apart. The purpose of most statistical models is to use imperfect data to provide an estimate of a population-level target of interest and the associated uncertainty of that estimate (from which statistical inference, e.g. confidence intervals or hypothesis tests can be calculated and performed). Leveraging spatial correlation when it exists can result in more precise estimates and ignoring it when it exists leads to incorrect (typically overly confident) uncertainty estimates (*22*). Non-probabilistic interpolation techniques cannot account for the sampling variability or produce reliable uncertainty estimates. In this work, we demonstrate a spatial statistical model that can be used to estimate biological fields across the tissue from which the spatialomics data sample was collected. The estimated fields represent the result that would be expected if the experiment were repeated many times.

One of the primary decisions to make when performing a spatial statistical analysis is whether the spatial domain (and thus the model) should be discrete or continuous in nature. Despite that many of the contemporary spatialomics data collection tools generate data on discrete “spot” arrays, the underlying tissue- and cellular-level *processes* — the target of our study — are not constrained to occur only at data observation locations (spots) or any finite set of locations.

For this reason, we have chosen to pursue a continuous spatial modeling approach. This does not mean that we believe there to be continuous distributions of discrete morphogen molecules at all points in space – we know this is not true – but rather that our estimate shows at each point in space the average values, and that spread of the values, that we would expect to see if we could repeat the experiment (of performing spatialomics data collection on a tissue slice) many times. Our goal with the spatial statistical models is two-fold: 1) to estimate the unknown underlying population field while 2) acknowledging the uncertainty in our estimate due to the sampling variability present in the observed dataset.

#### 2.1.3. A General Approach to Spatial Hierarchical Modeling

We formulate our model in a generalizable Bayesian hierarchical modeling framework that can generally be thought of as having three stages:

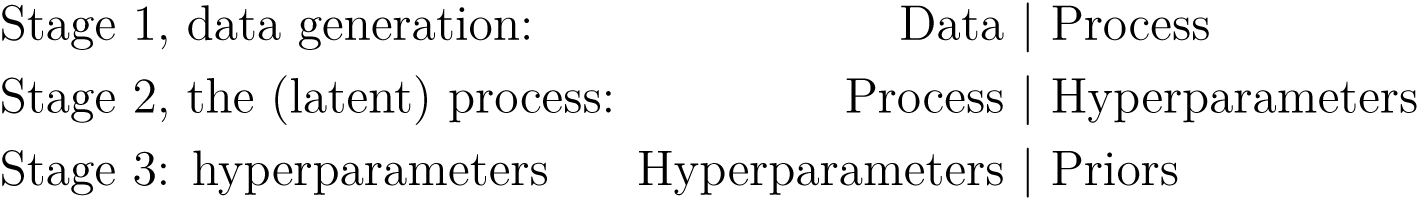

This broad framework is broadly applicable to many data collection settings where 1) data observations, which depend on some underlying process, are collected, 2) the underlying process, called the latent process since it cannot be directly observed, can be represented as a probability distribution whose parameters can be inferred from the data observations, and 3) the latent process itself may depend on some additional parameters. This simple yet powerful framework is well-established and a useful pedagogical tool for describing hierarchical modeling (*23*)”. To make this concrete in the spatialomics setting, we could define the following three stages as follows:

1. mRNA is collected across a tissue using a spatial transcriptomics data collection technique. At each location where data is collected, only a subset of all possible mRNA at that location is collected. Furthermore, the actual levels of mRNA at any particular location stochastically vary as mRNA is created and destroyed in accordance with the unobserved cell state at and around that location. The latent cellular state cannot directly be observed, but we aim to infer something about cell state using the observable mRNA counts.
2. The cell state, the latent process, is also a stochastic process. Conditional on the cell type, cell size, cellular microenvironment and signaling networks a cell is involved in, the state of the cell follows a probabilistic distribution. The state of the cell and the previously listed cell characteristics drive the stochastic mRNA production discussed in the above description of the data collection stage.
3. To complete the Bayesian specification, some of values that the cell state depends upon (e.g. the microenvironment of a cell) can also be viewed as stochastic processes whose parameters can be specified with prior distributions.

#### 2.1.4. A General Spatial Statistical Model for Spatialomics

The flexible and general purpose geostatistical model described in (1) can be applied to most, if not all, existing spatialomics datasets. Unlike many contemporary spatialomics and genomics data analyses which use some version of standardized (“normalized”) counts, we have proposed to directly model the “raw” counts from the spatialomics platform. This allows us to work with a generative model: the proposed model can be viewed as one capable of generating the data that is actually observed and collected when one uses a spatialomics platform.

Let ***s*** ∈ ***S*** denote a spatial location over continuous spatial domain ***S***. We will generally use *F* (***s***) to denote a continuously indexed spatial field, *F*, at location ***s***, and we denote the spatially varying distribution of the raw counts for the *i^th^* feature as *F_i_*(***s***).

To explicitly account for the differences between the raw counts and standardized counts, we specify the counts from feature *i* as the product of the total mRNA counts, N(***s***), and a rate parameter, *λ_i_*(***s***), which represents the expected counts of the *i^th^* feature per mRNA count at location ***s***. That is, *F_i_*(***s***) = N(***s***) × *λ_i_*(***s***).

The general hierarchical model then takes the form:

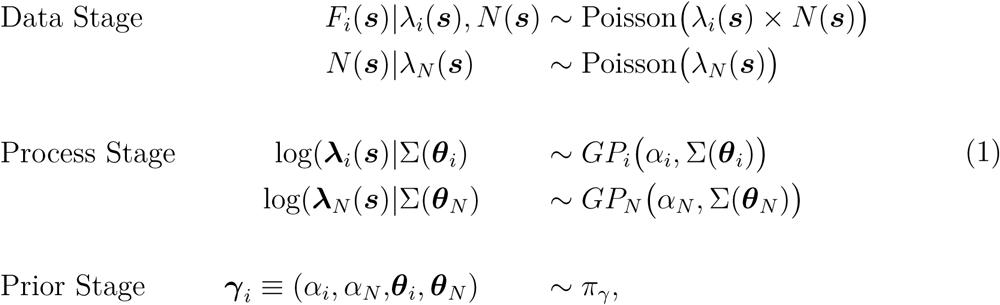

where we have modeled the counts of feature *i* at location ***s***, *F_i_*(***s***), as a Poisson distribution with rate equal to the total counts at ***s***, *N* (***s***), multiplied by the density of feature *i*, *λ_i_*, which is log-linked to a linear predictor consisting of a spatially structured Gaussian Process, *GP_i_* with Matérn covariance and mean *α_i_*. The nUMI, *N* (***s***), is similarly modeled. The Bayesian model specification is completed by specifying the hyperprior, *π_γ_*, over all all hyperparameters, ***γ****_i_*. Specifically, we set independent wide Gaussian priors on *α_i_* and *α_N_*, and we specified a joint penalized complexity prior on the range and variance parameters of each of the two Matérn covariance (*24*).

Due to the large number zeros often present in transcriptomics datasets, we also tested variations on the modeling framework wherein we replaced the Poisson likelihood for the *i^th^* feature with either a zero-inflated Poisson (ZIP) process or a zero-adjusted Poisson (ZAP) process. Both of these alternatives allow for a spatially varying probability of observing no counts. In these variants, the data generating mechanism can then viewed as first flipping a coin to determine if any counts from feature *i* will be observed, followed by a data observation from a Poisson process if the coin flip succeeds. Additional detail can be found in Section 2.2.4.

#### 2.1.5. Quantification of Categorical Spatial Structure

To assess the impact of our modeling choices, we desired a way of quantifying the spatial coherence of our final output clusters, represented as a categorical variable. To this end, we employed ELSA (entropy-based local indicator of spatial association), which is applicable to categorical variables and does not depend on the number of categories (*25*). ELSA is calculated by multiplying a spatially localized (scaled) Shannon entropy calculation with a localized measure of spatial dissimilarity to produce a measure of spatial association. Each of these components are bounded between 0 and 1, ensuring that ELSA is also bounded between 0 and 1. Locations with high values of ELSA indicate spatial regions where the variable of interest is less structured and spatially disparate while low values of ELSA suggest higher levels spatial coherence within the local neighborhood.

### 2.2. Computational Experiments

#### 2.2.1. Dataset Selection

To demonstrate our approach in detail, we leverage data from Chen et al. 2022 (*26*), a spatial time-course atlas of mouse embryogenesis acquired using the spatial transcriptomics platform Stereo-seq. This dataset is beautiful, information-rich, easy to interpret, and highly relevant to both developmental and regenerative biology. We selected a single field of view from one of the Day 16 timepoints, allowing close study of key morphogenic field interactions during development.

#### 2.2.2. Dataset Preprocessing

Raw count data were loaded into R and a subregion of the embryonic tissue was chosen to be visually recognizable by both experts and non-specialists. We then filtered the transcriptomic features to only those measured within this boundary, and, following the workflow from the original publication, collapsed the ultra-high resolution, sparse data to different sized spatial bins (10, 25, and 50). All the results shown in the figures of this manuscript use the bin-50 data.

To identify ligand and receptor genes, we cross-referenced the mouse version of the FANTOM5 database (*27*) against the feature names of the binned data, yielding 974 distinct genes that were either ligands or receptors. We further refined this gene list to only include ligands or receptors showing at least two counts in at least one cell, and at least one-quarter-of-one-percent of the total cells in the field of view expressing the gene at any non-zero level (see Supplement 2 for the visual outputs which were used to decide this threshold). This final step of feature selection yielded 918 individual ligand or receptor genes for modeling, which, based on the pairings in the FANTOM5 data base, would allow downstream computation of 1481 unique ligand-receptor interaction fields.

#### 2.2.3. Dataset Modeling

We individually modeled the spatial fields of the counts of the 918 features using a spatial Gaussian process model with Poisson observations, as defined in (1). The model was fit using R-INLA and predictions from these models were then used in downstream analyses.

#### 2.2.4. Comparison of 3 Data Models

We experimented with three different likelihood models: a Poisson likelihood, a zero-inflated Poisson (ZIP) likelihood, and a zero-adjusted Poisson likelihood (ZAP). The ZIP and ZAP models provide extra flexibility to acknowledge that the raw count data is often very sparse, with many spots returning a measurement of 0 for any particular feature:

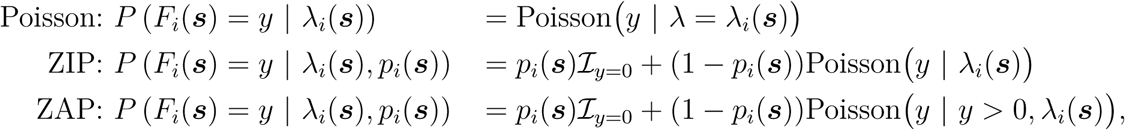

where *p_i_*(***s***) is a spatially varying probability of a 0 observation for feature *i*, which is modeled with a spatial Gaussian random field using a logit-link, and I*_y_*_=0_ is an indicator function which takes a value of 1 if *y* = 0 and takes a value of 0 otherwise. The ZIP and ZAP models differ in that the probability of a 0 in the ZAP model is directly governed by *p_i_*(***s***)) and allows for the probability of a 0 to be less than a standard Poisson, whereas the ZIP models can only increase the probability of a 0. ZAP models are typically selected when the process for a 0 occurrence differs substantially from the process driving the non-0 values.

At each observed location, ***s****_j_*, model performance was assessed using three distinct prediction scores combining the observed counts for feature *i* at ***s****_j_*, *y_i_*(***s****_j_*) and the features of the posterior predictive distribution of *F_i_*(***s***):

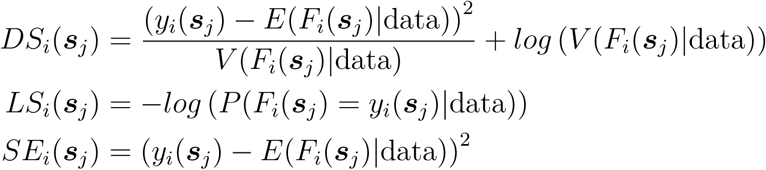

The Dawid-Sebastiani (DS) score is a proper scoring rule combining the predictive mean and variance, *E*(*F_i_*(***s***)|data) and *V* (*F_i_*(***s***)|data), respectively. The squared error (SE) is a proper score for the expectation, and the (negated) Log score (LS) is a strictly proper score (*28*).

#### 2.2.5. Quantitative Optimization of Ligand-Receptor Convolution

We experimented with multiple different approaches to ligand-receptor convolution. In all approaches studied, we did not consider ligand nor receptor sequestration due to binding, nor did we consider cross-mechanism inhibition or competitive interference, which can be the focus of future studies. We tested product, geometric mean, and a new approach labeled ‘common minimum convolution’, which computed the maximum possible number of local binding events, assuming that all ligand and all receptor bind to one another in a 1:1 fashion. We also explored a prototype approach based on chemical kinetics, using the rate equation with a tunable dissociation constant to model local steady-state ratio between bound and unbound ligand.

Our goal was to learn which methods yielded the most fine-grained and spatially organized clusters by looking for methods that maximized the number of clusters while minimizing the spatial entropy as quantified by ELSA (see Section 2.1.5). We therefore performed a targeted optimization experiment to identify which combination of parameters, models, and data inputs, all else being equal, yielded an output that maximized our unbiased identification of distinct local microenvironments while minimizing inter-cluster spatial entropy. ELSA was calculated locally and averaged across the modeling region to produce a single comprehensive score. This provided a means to visualize the boundaries between clusters, assessing how well they segregate in space, and to rank each computational approach by both number of clusters identified (all other parameters being equal) and the global spatial entropy of cluster segregation.

#### 2.2.6. Quantifying the Relationship Between Spatial Structure and Hyperfield Dimensionality

We used the results from a specific optimized workflow (Raw Counts - Poisson Model - Common Minimum Convolution) to quantify the full relationship between spatial domain architecture and signaling hyperfield dimensionality, because contemporary biologic theory posits that the complex spatial domains observed in tissues are likely a multivariable function of high-dimensional signal combinations rather than of a small set of select mechanisms. We therefore performed an experiment in which we iteratively increased the number of individual features considered for latent-space embedding and clustering and observed standardized unbiased outputs. Individual ligand-receptor mechanism features were first ranked based on system-scale variance, and then a for-loop was written to embed and cluster the data, with otherwise fixed algorithmic parameters, using increasingly many ligand-receptor features (10, 50, 100, 150, …, 1481). Ligand-receptor connectivity field data were scaled (though, importantly, not sum-normalized first) and processed without any principal-component analysis. Latent-space neighborhood graph construction and clustering parameters were held constant to allow isolation of the effect of feature number. ELSA was run in all instances, with a scan across the local distance parameter. All outputs from each step were stored for downstream visualization and analysis.

## 3. Results

### 3.1. Raw counts are optimal input for spatial modeling

Our experiments first revealed that commonly-applied count-normalization approaches create significant topological artifacts when applied to spatial data, a pattern also noted by others (*29*). Low individual-feature counts can be transformed into local hotspots when the raw values are count-normalized within spots of low total RNA density. See Figure 3 for a demonstration with a single feature (Cd44); similar comparisons between raw data vs. count-normalization for all features studied can be observed in Supplement 3. Because we aimed to model raw ligand-receptor *binding density* within tissues, the results from this experiment made clear to us that it was most physically appropriate to convolve continuous ligand and receptor fields which estimate *raw* ligand and receptor topology, rather than fields which estimate the commonly-used *count-normalized*topology. Nevertheless, we decided to carry forward both approaches in parallel experimentally, so that we could identify and compare project-relevant differences downstream. As a secondary result, we also learned form these early experiments that spatially modeling *N* (***s***), the nUMI, in addition to a ligand or receptor feature, *F_i_*(***s***), didn’t practically alter the outputs, and the remainder of the results were run using the nUMI directly, as measured in the data.

**Figure 1:**
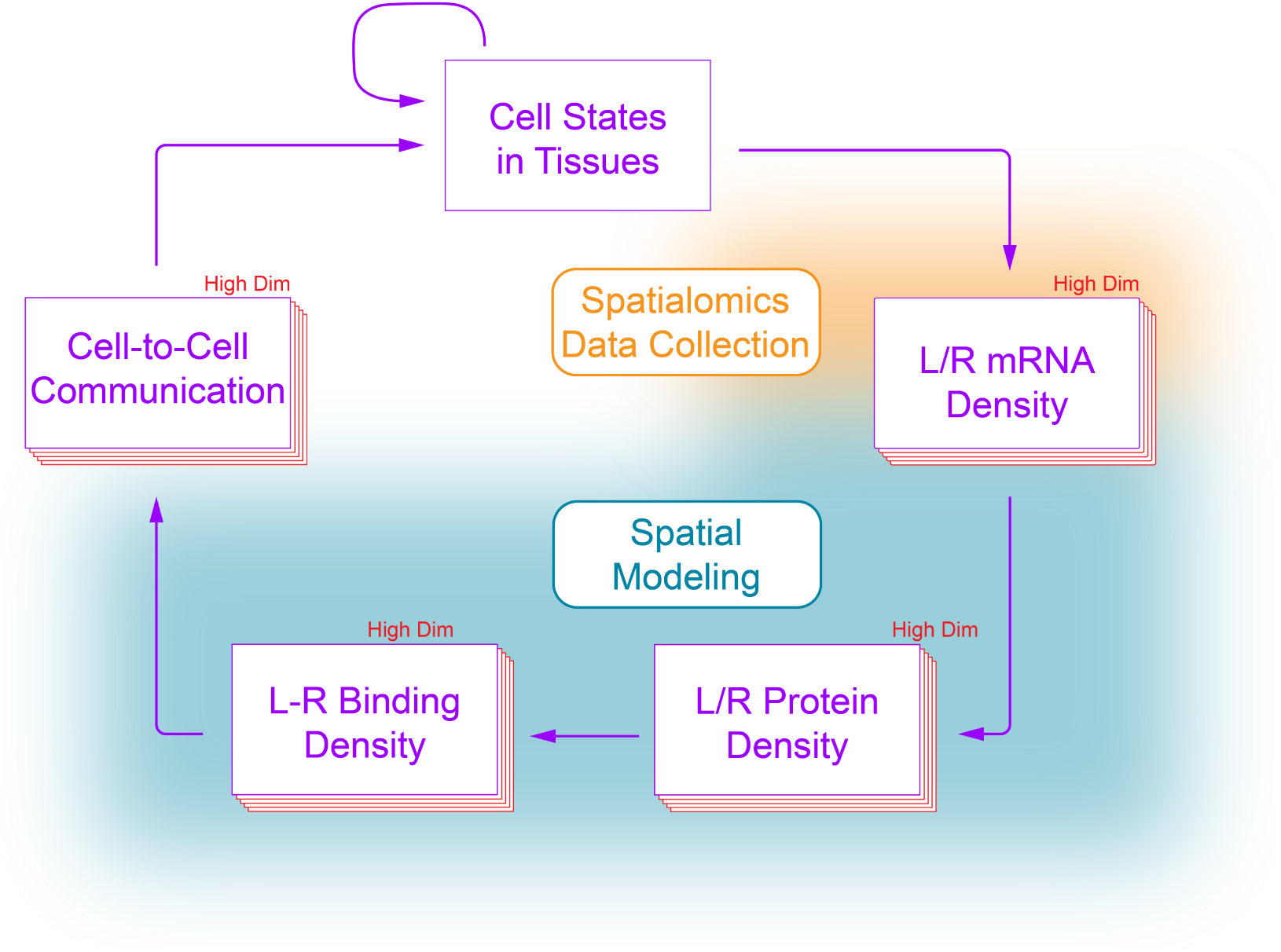
The purple arrows represent true tissue processing and activity. The blurred regions represent coarse estimates captured by spatialomics or created by spatial modeling.

**Figure 2:**
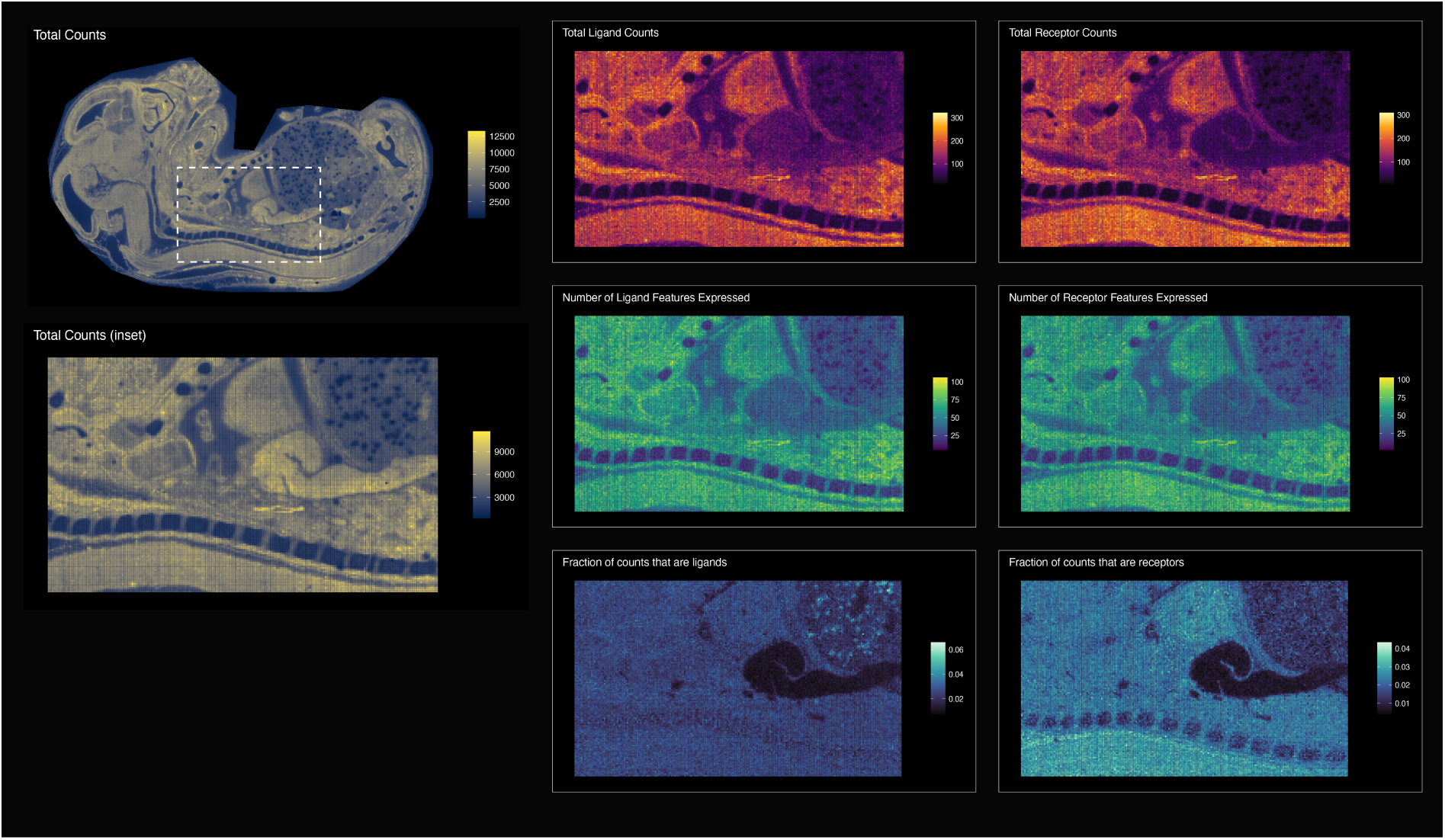
The 1^st^ columns shows total counts within the selected field of view from (*26*). The 2^nd^ column shows ligand information, and the 3^rd^ column shows receptor information. For latter two columns, the 1^st^ row is total ligand or receptor counts, the 2^nd^ row is total ligand or receptor features, and the 3^rd^ row is fraction of the total counts that are ligand or receptor counts.

**Figure 3:**
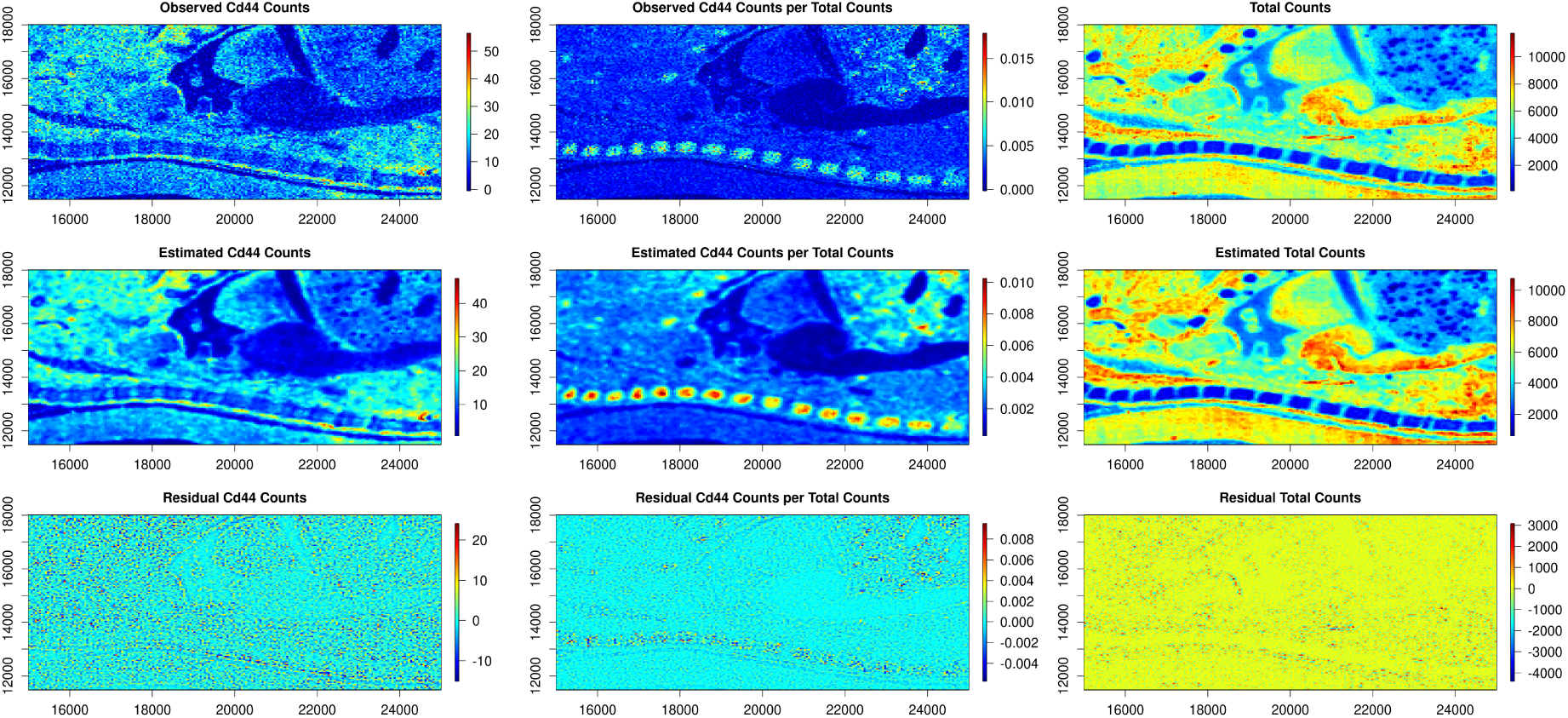
Example comparison between raw count topology vs. count-normalized topology for a single feature (Cd44). 1^st^ column is observed Cd44 counts, 2^nd^ column is observed Cd44 counts divided by the total counts. 3^rd^ column is the total counts. 1^st^ row is raw data, 2^nd^ row are estimates from the spatial model, 3^rd^ row is the residual (difference between model and raw data.) Count-normalization causes significant changes to spatial topology; we have concluded that normalization should be avoided prior to connectivity computations.

### 3.2. Poisson likelihood model outperforms ZIP and ZAP models

We compared three main spatial modeling approaches: POI, ZIP, and ZAP (see Methods). We found that the relative simplicity of the spatial Poisson model proved most suited, in these particular exercises, to the general task of estimating feature counts. Table 1 provides the average mean absolute errors, root mean square errors, Dawid-Sebastiani scores, and log scores, split out by likelihood and averaged over all observations and all 918 features. Predictive plots showed that the empirical distribution of the PIT scores from the Poisson models was routinely closest to a uniform distribution when compared to the empirical CDF of the PIT scores from the ZIP and ZAP models. Spatial residual plots revealed that the poorer average performance by the ZIP and ZAP models appears attributable to oversmoothing as evidenced by the predictions and residuals. See Figure 4 for comparison between models for a single feature. See Supplement 2 for corresponding plots for all features.

**Figure 4:**
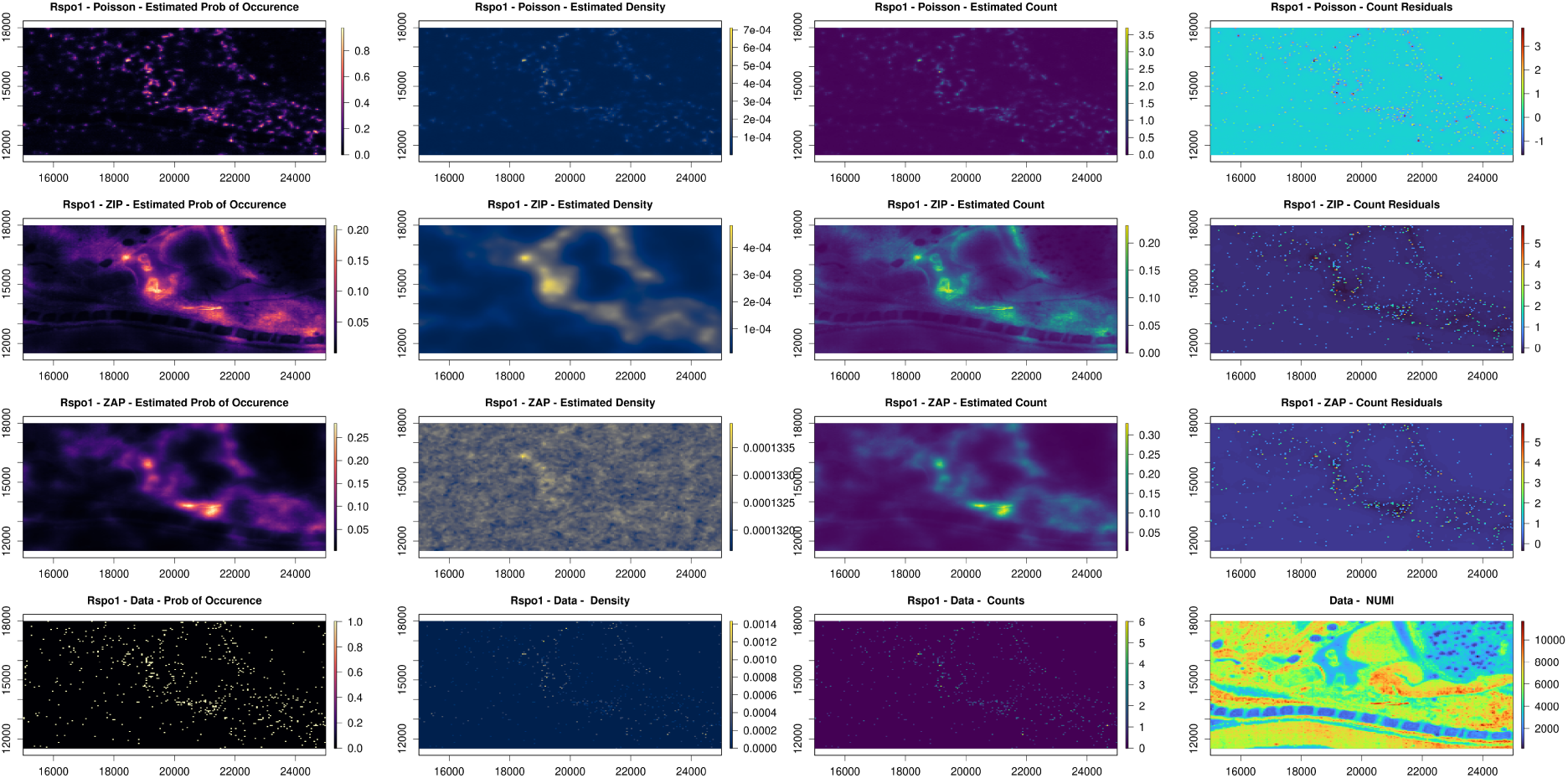
The 1^st^, 2^nd^, and 3^rd^ rows, respectively, show the outputs from the Poisson, ZIP, and ZAP models. The 4^th^ row shows raw data. The 1^st^ column shows the probability of a non-zero value at that point in space. The 2^nd^ column shows the density for that feature, i.e., counts/nUMI. The third column shows the feature counts. The top three rows in the 4^th^ column show the residuals, the difference between estimated counts and data counts, and the bottom right panel shows the nUMI from the data. The POI model resulted in the smallest residuals and was the best performing spatial model tested.

**Table 1:** Mean Absolute Errors, Root Mean Square Errors, Dawid-Sebastiani Scores, and Log Scores, by likelihood, averaged over all observations and all 1481 modeled transcriptomic features. The average percentage difference in the values, relative to the performance of the Poisson model, is given in parentheses.

| Model | MAE | RMSE | MDS | MLG |
| --- | --- | --- | --- | --- |
| Poisson | 0.21 | 0.42 | -2.15 | 0.31 |
| ZIP | 0.29 (111%) | 0.54 (116%) | -1.78 (85%) | 0.33 (111%) |
| ZAP | 0.23 (111%) | 0.47 (116%) | -1.77 (82%) | 0.35 (122%) |

### 3.3. Common minimum L-R convolution maximizes spatial structure

We assessed the output spatial structure that resulted from clustering 32 distinct spatial assays, summarized in Supplement [3]. Three in particular are detailed in Figure 5. We found that the ‘common minimum’ approach, which simply takes the common minimum between the ligand and receptor subunits, maximizes the number of clusters identifiable in the data while simultaneously minimizing spatial entropy. Of note, we found that the common minimum approach closely approximates the binding density outputs which are produced when *k*, the dissociation constant, is decreased to 0. This corresponds to total binding without any dissociation. Because of these results, we determined this to be an optimal output of our model to use for downstream experimentation. For full results from this experiment, see Supplement [4].

**Figure 5:**
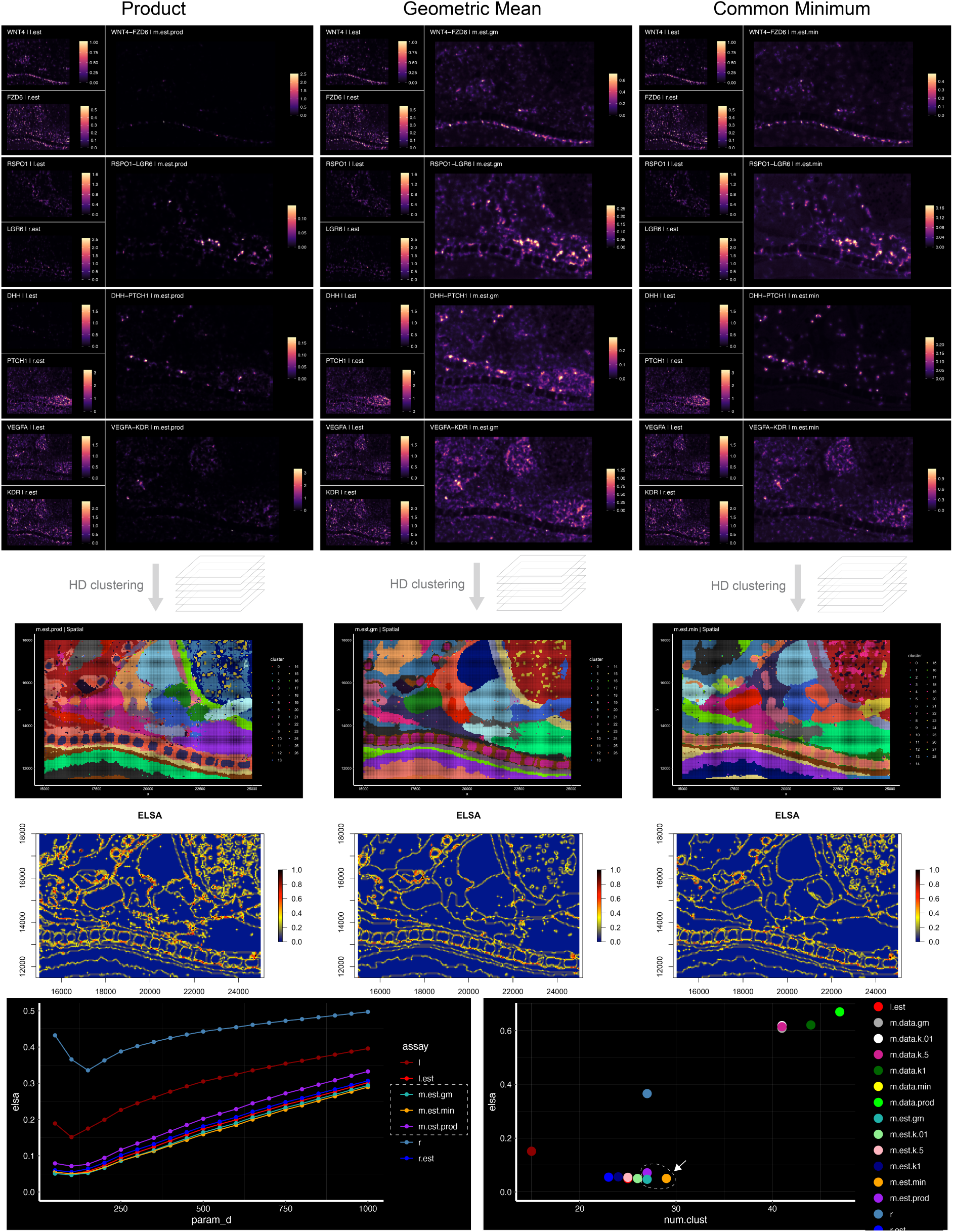
Comparison of convolution methods. Product, geometric mean, and common minimum shown in 1st, 2nd, and 3rd column, respectively. The values are less important than the relative topology, because the values are generally scaled for use in high-dimensional analysis. Middle of the page shows high dimensional clustering with a fixed workflow (see Methods). Cluster color and number is arbitrary; cluster number and spatial organization is relevant. ‘Entrograms’, made using ELSA (*30*), quantify local spatial coherence. The three methods detailed here are the top performing, maximizing cluster number while minimizing ELSA. The common minimum approach (‘m.est.min’) is the top performing of all methods tested.

### 3.4. High-dimensional morphogenic field analysis resolves spatial domains

To test our central hypothesis, that tissue spatial domains are a direct function of signaling field hyperstructure, we performed a targeted parameter sweep experiment in which we iteratively increased the number of signaling features, ranking them by variance, while holding all else equal. We iteratively added more features to our analysis and quantified the output number of clusters and their spatial entropy to observe changes in spatial domain architecture in both UMAP and histologic space. Our results are summarized in Figure 6, and full results may be found in Supplement ***[5]

**Figure 6:**
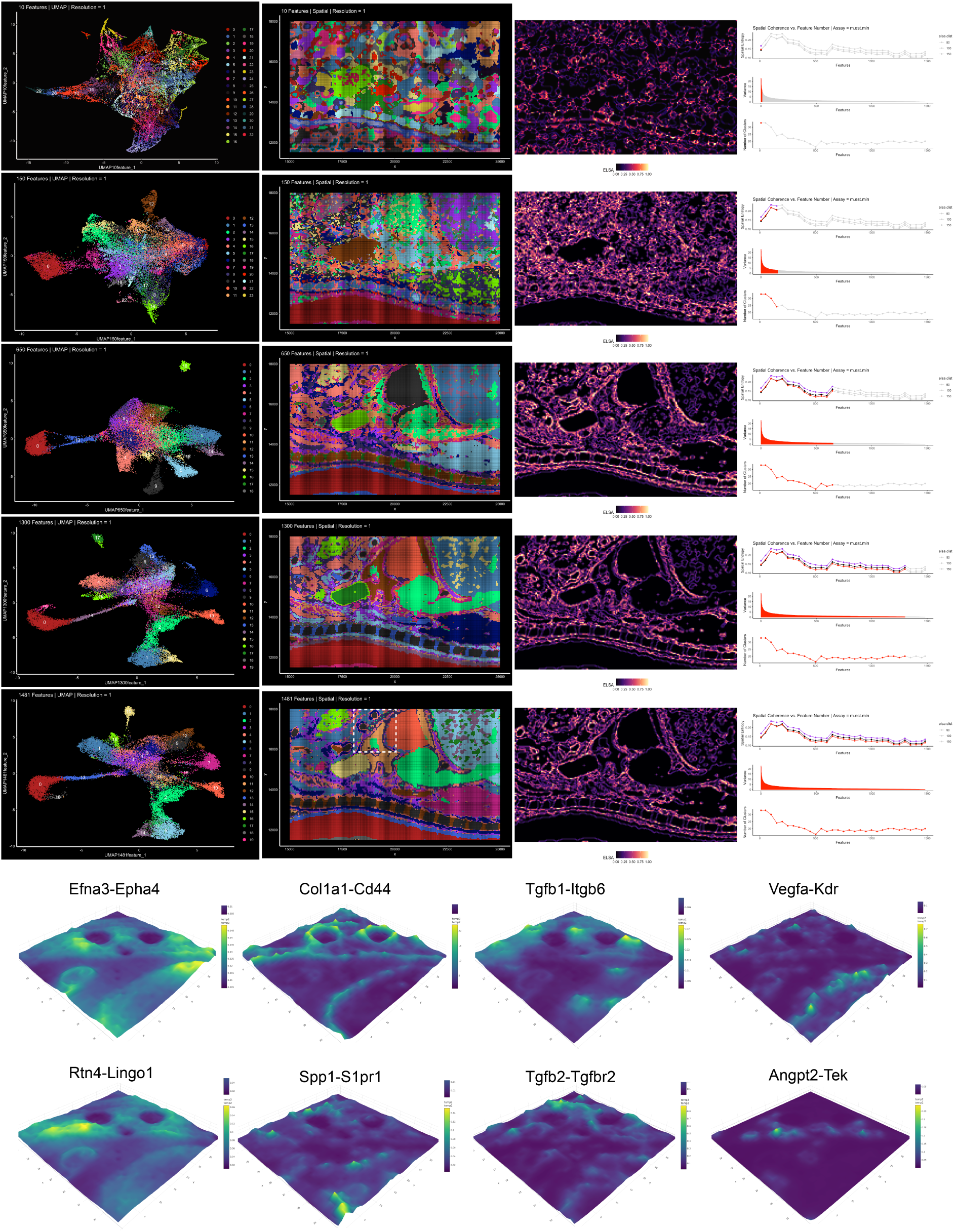
Top five rows show increasing spatial domain definition and organization as the number of morphogenic interaction fields are increased. Endpoint spatial domain architecture is a detailed function of all 1,481 unique signaling mechanisms, treated without principle-component analysis (see Methods). Bottom two rows show 8 selected morphogenic interaction fields of biologic interest, represented as three-dimensional density plots, corresponding to the tissue region within the dotted white line.

We found the spatial domain architecture in tissues to be a high-dimensional function of many signaling mechanisms at once. The effect is non-linear, with individual feature variance not directly predicting of the effect of adding said feature into holistic analysis. This finding supported the non-reducibility of hyperfield information. We found that clustering the data without any principal-component analysis resulted in the greatest spatial complexity and coherence. Clustering the tissue within the full 1,481-dimension feature-space maximized spatial complexity and coherence, allowing the resolving of subtle local detail, such as inter-organ fascial planes, that was unobserved when all features were not included or when dimensional-reduction was employed via PCA.

We closely observed hyperfield structure in a small sub-region of the embryo, outlined by a dotted-white boundary in 6. Selected feature-level topologies (individual lignad-recptor binding density field) have been rendered as three-dimensional surfaces at the bottom of 6. HTML files for all 1,481 fields within this region, allowing user-interactive zoom, value estimation and rotation in space, are available in Supplement ***[6].

## 4. Discussion

A central question in tissue biology is how and why tissues develop, retain, and lose their form. This broad question is fundamental to developmental biology, tissue engineering, regenerative medicine, and multicellular biology across species. For the better part of a century, we have understood that extracellular signaling plays a critical role in the patterning and organization of cells within tissues. Local intercellular cues preserve adult stem cell niches, govern cellular differentiation, and regulate tissue response to injury. However, despite this knowledge, our ability to *understand* cellular communication in tissues, i.e., to deliberately influence it to induce healing or desired change, is woefully limited.

Recent research has revealed that the grammar of cellular communication is intrinsically high-dimensional, with cells communicating on hundreds-to-thousands of wavelengths at once and processing/interpreting these cues via whole-cell parallel processing (*7*). This means that in order to properly model and visualize cell-to-cell communication, we must be comfortable observing and interpreting high-dimensional signaling ‘hyper-fields’ as single, holistic objects of study in histologic space. The work in this paper has been carried out with this principle in mind.

We see the outputs from these models as the start of a growing foundation for what we envision as ‘morphogenic field analysis’ — the interdisciplinary study of how morphogenic field interactions, gradients, dynamics, and perturbations shape tissue organization and control cellular differentiation, ideally in three-dimensions over time. Although the data analysis shown here is exclusively two-dimensional, and does not include a time dimension, there are many datasets emerging which do allow spatiotemporal analysis and there are incoming tools allowing affordable 3D profiling (*31*). Our approach is easily adapted to 3-dimensional timeseries data. When applied to timeseries datasets, longitudinal quantitative modeling could be performed to show how tissue field dynamics precede, align with, or follow cellular growth and change during tissue remodeling. Our methods are generally-applicable to any spatial transcriptomic data when the right choices are made: spot-based vs. imaging-based transcriptomics, for instance, should be treated differently, as their measurement methods and organization are fundamentally different. Further, our model is easily customized for specific tissue types: we are currently experimenting with an approach optimized for lung tissue that allows exclusion of large lumens and alveolar airspaces from the tissue fields.

The statistical models presented here have some limitations. First, they must be tuned to each input type of data — Xenium, Visium, StereoSeq (shown here), CurioTrekker, CosMx, COMET, etc. However, our general approach has thus far proved easy to adapt on all data types tested. Second, we currently are ignoring three major considerations: 1. each signaling mechanism is treated the same (there is no incorporation of molecule-specific diffusion coefficients based on molecular weight), 2. there is no accommodation for variable extracellular matrix properties (competitive binding, protein-density-diffusion limitations), and 3. there is no accommodation for ligand-receptor sequestration due to binding (complex non-linear modeling required). These deficits shall be addressed in future studies, and present many interesting opportunities for further experimentation.

“All models are wrong, but some are useful”. The modeling outputs shown here are a statistically-powerful *proxy* for high-dimensional information flow *potential* within tissues. Further data must be acquired at the protein level and intracellular level before any of these model outputs are treated as evidence of actual information transmission or trans-duction. The experimental cost of doing this, however, can be high, particular if the patterns of interest are high-dimensional and cannot be reduced to targeted cell-scale, pathway-scale, or molecule-scale experiments. Therefore, in our opinion, the primary power of the modeling approach described in this paper lies not in its ability to predict single-mechanism interactions. Rather, it lies in its ability to facilitate statistically-robust mappings between holistic high-dimensional extracellular signaling milieus and cell-state-specific intracellular state changes and transitions. A cell within a tissue experiences hundreds to thousands of cues at once, and processes these cues in parallel — as a single, holistic, high-dimensional signal. Our modeling approach allows us to locally estimate this high-dimensional signal while providing measures of statistical confidence. When paired with *intra*cellular data, such as that regarding transcription factor activation, cytoskeletal remodeling, metabolic reprogramming and mechanoresponse, we are newly provided with an opportunity to quantitatively study the precise ways that different cells, in different states, process and respond to high-dimensional dynamics in their local extracellular milieu: i.e., how holistic cellular microenvironment governs cellular state — and therefore how extracellular milieus might be targeted in the future to induce regenerative healing.

Our code is open access on GitHub, and we have included all of our work from this project so that others can replicate our results, see how we have organized our inputs/outputs, and experiment on their own. The images presented in the figures of this paper are a small fraction of our model outputs — and we encourage interested readers to download the supplemental materials included with this paper as a way to observe the true scale and structure of extracellular communication in mammalian tissues. We look forward to working with the community to explore morphogenic field analysis in additional tissue datasets and to brainstorm ways to influence spatial extracellular signaling for therapeutic means.

## 5. Code Availability

All code used to create this work is available on GitHub at: https://github.com/aaron-oz/a-s-omics

## 6. Data Availability

Raw data used to create the figures in this manuscript can be downloaded from (*26*): https://db.cngb.org/data_resources/project/CNP0001543

## 7. Supplemental Materials

We have provided a set of documents on FigShare which capture key intermediate- and end-outputs of the modeling described in this paper. They can be downloaded from the following links:

Supplement 1 Raw-to-Bin: https://figshare.com/s/e921e8e36296cd8c57da
Supplement 2 Feature Refining: https://figshare.com/s/6656c05d4552a59afa05
Supplement 3 Raw vs. Norm, Selected Features: https://figshare.com/s/4885d58ba45b2f2e7689
Supplement 4 Model Comparison All Features: https://figshare.com/s/9d266d5f0fd39632986d
Supplement 5 HD Embeddings: https://figshare.com/s/7bbe3feb729a9731acfa
Supplement 6 ELSA Analysis: https://figshare.com/s/4867799e0e8b68fba5a8
Supplement 7 Clusters vs. Features: https://figshare.com/s/2e305ce9ee20efd7ee47
Supplement 8 Morphogen Fields: https://figshare.com/s/2f66def6d22f1549b312

## 8. Acknowledgments

This work was supported by T32GM086287 from the NIGMS, and laboratory startup funds from the Yale School of Medicine, the Yale Department of Anesthesiology, and Bucknell University. The opinions expressed are those of the authors and do not necessarily represent the thoughts or opinions of NIGMS, NIH, or the United States government.

